# SOS2 regulates the threshold of mutant *EGFR*-dependent oncogenesis

**DOI:** 10.1101/2023.01.20.524989

**Authors:** Patricia L. Theard, Amanda J. Linke, Nancy E. Sealover, Brianna R. Daley, Johnny Yang, Katherine Cox, Robert L Kortum

**Affiliations:** Department of Pharmacology and Molecular Therapeutics, Uniformed Services University of the Health Sciences, Bethesda, MD, USA 20814

**Author notes:** Competing Interests statement: The authors declare no competing financial interests.

## Abstract

Son of Sevenless 1 and 2 (SOS1 and SOS2) are RAS guanine nucleotide exchange factors (RasGEFs) that mediate physiologic and pathologic RTK-dependent RAS activation. Here, we show that SOS2 modulates the threshold of epidermal growth factor receptor (EGFR) signaling to regulate the efficacy of and resistance to the EGFR-TKI osimertinib in lung adenocarcinoma (LUAD). *SOS2* deletion sensitized *EGFR*-mutated cells to perturbations in EGFR signaling caused by reduced serum and/or osimertinib treatment to inhibit PI3K/AKT pathway activation, oncogenic transformation, and survival. Bypass RTK reactivation of PI3K/AKT signaling represents a common resistance mechanism to EGFR-TKIs; *SOS2* KO reduced PI3K/AKT reactivation to limit osimertinib resistance. In a forced HGF/MET-driven bypass model, *SOS2* KO inhibited HGF-stimulated PI3K signaling to block HGF-driven osimertinib resistance. Using a long term *in situ* resistance assay, a majority of osimertinib resistant cultures exhibited a hybrid epithelial/mesenchymal phenotype associated with reactivated RTK/AKT signaling. In contrast, RTK/AKT-dependent osimertinib resistance was markedly reduced by *SOS2* deletion; the few *SOS2* KO cultures that became osimertinib resistant primarily underwent non-RTK dependent EMT. Since bypass RTK reactivation and/or tertiary *EGFR* mutations represent the majority of osimertinib-resistant cancers, these data suggest that targeting SOS2 has the potential to eliminate the majority of osimertinib resistance.

**One sentence summary:** SOS2 modulates the threshold of EGFR-PI3K signaling to regulate the efficacy of and resistance to osimertinib.

## Introduction

Lung cancer is the leading cause of cancer death; lung adenocarcinoma (LUAD) is the most common subtype of lung cancer^1^. LUAD is primarily a disease of hyperactivated receptor tyrosine kinase (RTK)/RAS signaling, and 75-90% of LUADs harbor oncogenic driver mutations in RTK/RAS pathway members^2–4^. Activating epidermal growth factor receptor (EGFR) mutations drive oncogenesis in 15-30% of LUADs and are the major cause of LUAD in never-smokers^1^. For patients with *EGFR-*mutated LUAD, 1^st^ (gefitinib and erlotinib), 2^nd^ (afatinib and dacomitinib), and 3^rd^ (osimertinib) generation EGFR-TKIs (tyrosine kinase inhibitors) have revolutionized cancer treatment. However, despite markedly enhancing survival of patients with *EGFR*-mutant tumors, resistance to EGFR-TKIs invariably emerges. For 1^st^-generation EGFR-TKIs gefitinib and erlotinib, resistance is primarily driven by either mutations in the drug binding site (T790M; 60%) or oncogenic shift to alternative RTKs (15-30%). The 3^rd^-generation EGFR-TKI osimertinib was developed to target T790M-mutated EGFR; osimertinib showed enhanced progression-free^5^ and overall survival^6^ compared to 1^st^- and 2^nd^-generation EGFR-TKIs and is now the first-line treatment in *EGFR*-mutated LUAD. However, despite the increased effectiveness of osimertinib, resistance invariably emerges.

Similar to 1^st^-generation EGFR-TKI resistance, osimertinib resistance can be driven by both EGFR-dependent and EGFR-independent mechanisms, however, unlike 1^st^-generation EGFR-TKIs EGFR-independent mechanisms predominate^7–10^. While the frequency and types of resistance may depend on whether osimertinib was used as first-line therapy or second-line therapy after a patient developed resistance to 1st-generation EGFR-TKIs, the most common EGFR-independent resistance mechanisms involve reactivation of the RTK/RAS/effector pathway ^10^ via enhanced signaling through parallel RTKs^7–16^. While individual RTK inhibitors may be beneficial in cancers whose resistance is driven by a specific RTK (MET, AXL, HER2/3, FGFR), broad inhibition of RTK signaling is likely required to enhance osimertinib efficacy and delay therapeutic resistance^7–16^. Alternatively, a subset of osimertinib resistant tumors acquire resistance through histologic transformation via either epithelial-to-mesenchymal transformation (EMT) or transition to small cell lung cancer (SCLC). EMT is a dynamic process by which epithelial cells acquire mesenchymal characteristics via changes both in gene transcription and post-translational regulatory mechanisms and is often characterized by the loss of E-Cadherin and an increase in Vimentin abundance^17^. EMT is a common feature in RTK/RAS pathway independent osimertinib resistance, and targeting EMT via the transcription factors TWIST1^18^ or Snail^19^ re-sensitizes osimertinib-resistant cells to osimertinib.

The RASGEFs Son of Sevenless 1 and 2 (SOS1 and SOS2) mediate RTK-stimulated RAS activation and represent common proximal RTK pathway intermediates whose inhibition has the potential to delay therapeutic resistance driven by RTK reactivation^4, 20^. Although SOS1 and SOS2 were previously considered poor candidates for therapeutic intervention due to their low oncogenic potential, recent studies showed that both SOS1 and SOS2 may be important therapeutic targets in *EGFR*- and *KRAS-* mutated cancers^21–27^, and the SOS1 inhibitor BI-1701963 is currently in Phase I trials for treating *KRAS*-mutated cancers both as a single agent and in combination with KRAS^G12C^ [NCT04975256] or MEK [NCT04111458] inhibitors. SOS1 is an important mediator of mutant EGFR-driven oncogenesis, and, moreover, the preclinical SOS1 inhibitor BAY-293 synergizes with osimertinib to limit survival of 3D spheroid cultured LUAD cells^22^.

Here we show that SOS2 modulates the threshold of EGFR signaling to regulate the efficacy of and resistance to osimertinib in *EGFR*-mutated LUAD cells. Using mouse embryonic fibroblasts (MEFs) expressing mutated EGFR proteins, we found that mutant EGFR-driven transformation was more sensitive to perturbations in the level of EGFR stimulation in *Sos2* KO cells compared to WT controls. *Sos2* KO cells showed reduced mutant EGFR-driven transformation that was inhibited by low levels of EGFR-TKI treatment and restored by exogenous EGF stimulation. We observed similar results in *EGFR*-mutated LUAD cells. 3D spheroid growth and survival were more sensitive perturbation of RTK signaling caused by reduced serum conditions and/or treatment with the 3^rd^ generation EGFR-TKI osimertinib in *SOS2* KO cells compared to non-targeting controls.

RTK pathway reactivation represents a common mechanism driving resistance to EGFR-TKIs including osimertinib^4, 7–16^, and RTK-dependent PI3K/AKT activation is a common hallmark of EGFR-TKI resistance^28, 29^. Using a forced HGF/MEK-driven bypass model, we found that *SOS2* KO limited HGF-stimulated AKT signaling and blocked HGF-driven recalcitrance to osimertinib therapy. Using long term *in situ* resistance assays (ISRAs)^30^, we found that a majority of osimertinib resistant cultures exhibited a hybrid epithelial/mesenchymal phenotype associated with reactivated RTK/AKT signaling. In contrast, the number of RTK/AKT-dependent resistant cultures was markedly reduced by *SOS2* deletion, with the few resistant *SOS2* KO cultures that did emerge doing so primarily by undergoing non-RTK dependent EMT. These data suggest that targeting SOS2 has the potential to eliminate the majority of osimertinib resistance, since bypass RTK reactivation and/or tertiary *EGFR* mutations represent the majority of osimertinib-resistant cancers.

## Results

### SOS1 and SOS2 mediate mutant EGFR-dependent transformation

To investigate the independent and combined roles of SOS1 and SOS2 in mutant EGFR-driven oncogenesis, we assessed anchorage-independent growth in immortalized WT, *Sos1*^-/-^, *Sos2*^-/-^, and *Sos1/2* DKO MEFs^21^ expressing WT or mutated EGFR (Fig. 1) in both the absence and presence of EGF stimulation. EGF stimulation was performed as a large proportion of lung adenocarcinomas show high expression of EGFR ligands^31^ and seminal experiments showed that EGF stimulation promoted transformation in cells overexpressing WT EGFR and enhanced transformation in cell expressing oncogenic EGFR mutants^32, 33^. In the absence of exogenous EGF stimulation, we found that in addition to SOS1, SOS2 was a critical modifier of mutant EGFR-driven transformation (Fig. 1, open bars). *Sos1* or *Sos2* deletion reduced mutant EGFR-driven transformation in the absence of exogenous EGF, and mutant EGFR-driven transformation was completely abrogated by combined *Sos1/2* KO. These data revealed a previously uncharacterized role for SOS2 in mutant EGFR-driven transformation.

**Fig 1.**
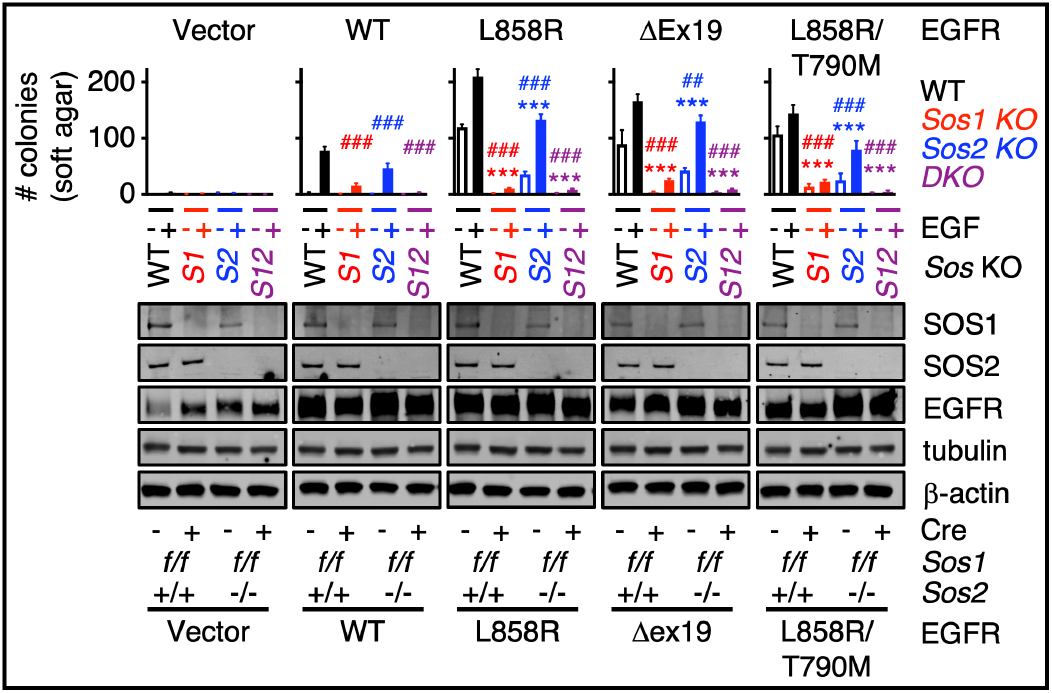
SOS1and SOS2 mediate mutant EGFR-dependent oncogenic transformation. *Sos1^f/f^* and *Sos1^f/f^Sos2^-/-^* MEFs were transduced with lentiviruses expressing either empty vector or the indicated WT or oncogenic EGFR mutants; *Sos1* was then deleted by retroviral transduction with Cre recombinase to generate WT (black) *Sos1^-/-^* (red), *Sos2^-/-^* (blue) and *Sos1^-/-^ Sos2^-/-^* (purple) MEFs. Cells were assessed for colony growth in soft agar to assess anchorage independent growth in the absence (open) or presence (filled) of exogenous EGF-stimulation (20 ng/mL) (top). Whole cell lysates (WCLs) were analyzed by Western blotting with antibodies specific for SOS1, SOS2, EGFR, tubulin, or β-actin (bottom). Statistical significance was determined by ANOVA using Tukey’s method for multiple comparisons from *N*=4 independent experiments. * p<0.05, **p<0.01, ***p<0.001 vs. unstimulated NT cells. # p<0.05, ## p<0.01, ### p<0.001 vs. *EGF-*stimulated NT cells. Data are from four independent experiments. Western blots are representative from *N*=4 independent experiments.

Upon EGF stimulation, MEFs overexpressing WT EGFR showed transforming growth and MEFs expressing mutated EGFR proteins showed a 1.5-2-fold increase in transformed colonies, confirming a role in ligand-dependent enhancement of EGFR-driven oncogenesis. Intriguingly, while EGF stimulation only modestly enhanced transformation in *Sos1*^-/-^ cells, EGF stimulation restored mutant EGFR-driven transformation in *Sos2*^-/-^ cells (Fig. 1 and 2A). These data suggest that SOS2 may modulate threshold of EGFR signaling required to promote oncogenesis.

**Fig 2.**
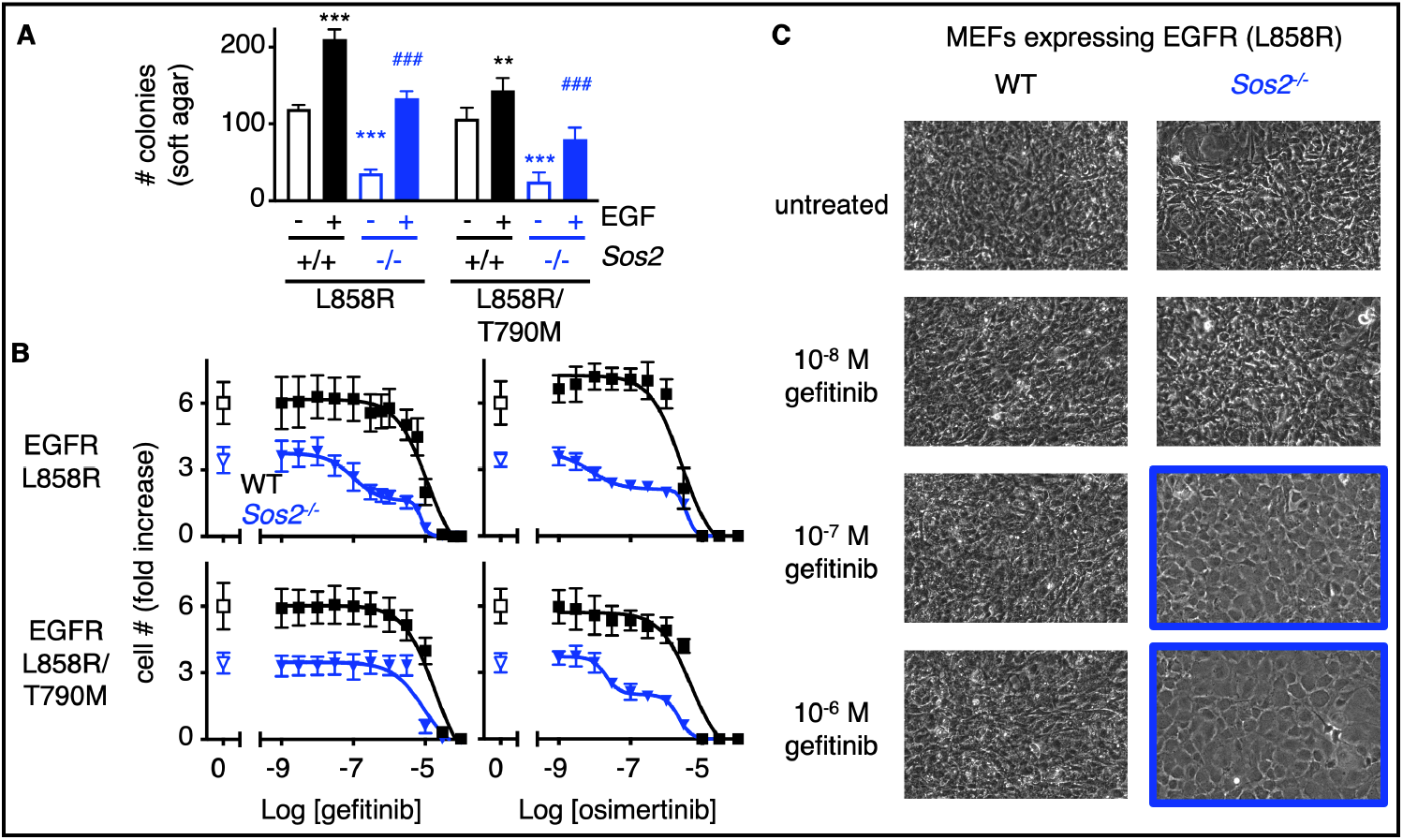
*Sos2* deletion synergizes with EGFR-TKI treatment to inhibit mutant EGFR-driven transformation. (**A**) Soft agar assays from WT and *Sos2^-/-^* MEFs expressing L858R or L858R/T790M mutated EGFR from Fig. 1. (**B**) Dose-response curves of WT (black squares) or *Sos2*^-/-^ (blue inverted triangles) MEFs expressing 1^st^ generation EGFR-TKI sensitive (L858R) or resistant (L858R/T790M) mutated EGFR from Fig. 1 treated with increasing doses of the 1^st^-generation EGFR-TKI gefitinib (left) or the 3^rd^-generation EGFR-TKI osimertinib (right) for five days. Dose-response curves are normalized to cell number assessed two hours after plating by CellTitre Glo. Data are presented as mean +/-s.d. from *N*=3 independent experiments. (**C**) 10 x photographs of post-confluent *Sos2^+/+^* or *Sos2^-/-^* MEFs expressing EGFR (L858R) treated with the indicated dose of gefitinib from (B). Data are representative from three independent experiments.

### SOS2 regulates the threshold of EGFR-TKI dependent inhibition of oncogenesis

SOS1 inhibition synergized with EGFR-TKIs to limit cell survival under short-term culture conditions^22^, however, whether SOS1 or SOS2 ablation enhance the ability of EGFR-TKIs to inhibit oncogenic transformation has not been assessed. To directly test the extent to which SOS1 and/or SOS2 modify EGFR-TKI-dependent inhibition of oncogenesis, we assessed dose-dependent changes in cell number in WT, *Sos1* KO, *Sos2* KO, and *Sos1*/2 DKO MEFs expressing both first-generation EGFR-TKI sensitive (L858R) and resistant (L858R/T790M) EGFR mutants. MEFs were seeded in 96-well cell culture plates and grown for 48 hours. Cells were approximately 50% confluent prior to treatment with increasing doses of the first-generation EGFR-TKI gefitinib or the third-generation EGFR TKI osimertinib for five days (Fig. 2B and S1). This cell density allows for the assessment of post-confluent cell growth due to loss of contact inhibition; untreated WT MEFs expressing mutant EGFR showed a roughly 6-fold increase in cell number over the five-day period. In WT MEFs expressing EGFR (L858R), gefitinib and erlotinib inhibited cell outgrowth at very high levels of drug (EC_50_ ∼ 10 μM), indicative of general toxicity rather than on-target inhibition. Although both the *Sos1* or combined *Sos1/2* deletion inhibited cell growth in the absence of EGFR-TKI treatment, neither altered the EC_50_ for either gefitinib or osimertinib (Fig. S1).

In contrast, *Sos2*^-/-^ cells expressing a first-generation EGFR-TKI sensitive mutant [EGFR (L858R)] showed a biphasic response to both gefitinib and osimertinib with the first inflection roughly 2-log lower than the toxic dose for either drug. *Sos2*^-/-^ cells expressing a first-generation EGFR-TKI resistant mutant [EGFR (L858R/T790M)] were unresponsive to gefitinib but showed a similar biphasic response to osimertinib treatment (Fig. 2c). To confirm that the first EGFR-TKI-dependent inhibition of cell number in *Sos2*^-/-^ cells was due to inhibiting transformation, WT and *Sos2*^-/-^ MEFs expressing EGFR (L858R) were treated with increasing doses of gefitinib for two weeks (1-week post-confluence) and transformation was assessed by loss-of-contact inhibition. WT MEFs showed loss-of-contact inhibition (transformation) at gefitinib doses up to 1μM. In contrast, *Sos2*^-/-^ MEFs treated with ≥ 100 nM gefitinib were contact inhibited and grew as a monolayer (Fig. 2C). These data suggest SOS2 may be an important modifier of oncogenic growth and EGFR-TKI responsiveness in *EGFR*-mutated cancer cells.

We next assessed the extent to which SOS1 or SOS2 regulate the threshold of EGFR signaling to promote oncogenesis in human *EGFR*-mutated LUAD cells. *SOS1* or *SOS2* were deleted in a panel of *EGFR*-mutated LUAD cell lines (Fig. 3A). 3D spheroid growth was assessed over seven (PC9) or 21 (H1975, HCC827) days at decreasing serum concentrations in either untreated cells (Fig. 3B) or at increasing osimertinib concentrations (Fig. 3C). For all CRISPR experiments, we assessed the effect of *SOS1* or *SOS2* deletion from cell populations that showed >80% decreases in SOS1 or SOS2 protein abundance compared to NT controls; populations were used rather than cell clones to avoid clonal effects not related to *SOS1* or *SOS2* KO. Compared to non-targeting (NT) controls, *SOS1* KO significantly reduced transformation in PC9 cells and completely inhibited transformation in H1975 and HCC827 cells in agreement with previous findings^22^. In contrast, the effect of *SOS2* deletion on transformation was serum-dependent. While *SOS2* KO had a modest effect on transformation in 10% serum, the dependence of transformation on SOS2 was more pronounced as serum concentrations decreased so that at low serum *SOS2* KO cells showed inhibition of transformation comparable to *SOS1* KO cells (Fig. 3B). These data suggest a critical role for SOS2 in mutant EGFR-driven transformation under nutrient limiting conditions.

**Fig. 3.**
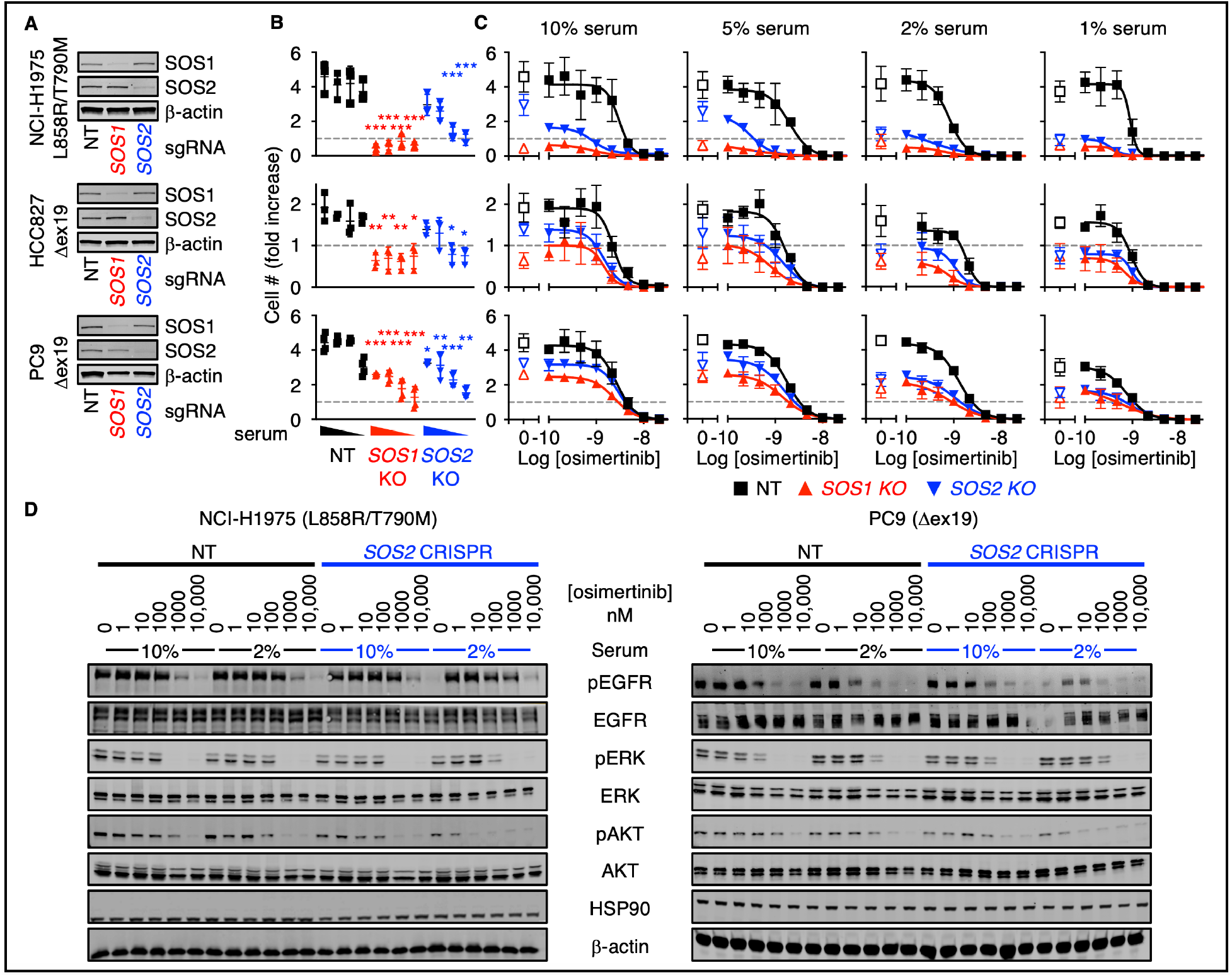
*SOS2* deletion increases the threshold of EGFR stimulated required for oncogenic transformation. (**A** to **B**) 3D spheroid growth under decreasing serum concentrations (10%, 5%, 2%, and 1%) in the absence of EGFR-TKI treatment (A) or at increasing doses of osimertinib in pooled populations of PC9, H1975, or HCC827 cells where *SOS1* or *SOS2* has been deleted using CRISPR/Cas9 versus NT controls after 7 (PC9) or 21 (H1975, HCC827) days to allow for transforming growth (B). (**C**) Western blots of WCLs showing *SOS1* or *SOS2* deletion. (**D**) Western blots WCLs from 3D spheroid cultured *SOS2* KO NCI-H1975 (left) or PC9 (right) cells versus NT controls treated with increasing doses of osimertinib under high serum (10%) or low serum (2%) conditions for 6 hours. Western blots are for pEGFR, EGFR, pERK, ERK, pAKT, AKT, HSP90, and β-actin. Data are presented as mean +/-s.d. from *N*=3 independent experiments. Western blots are representative of *N*=3 independent experiments.

We further assessed the extent to which *SOS1* or *SOS2* deletion enhanced osimertinib dose-dependent inhibition of transformation (above grey line, Fig. 3C) and survival (below grey line, Fig. 3C) in long-term 3D spheroid cultured LUAD cells. In NT controls, osimertinib caused a dose-dependent decrease in transforming growth at low doses and inhibited survival at higher doses. *SOS1* KO alone was sufficient to inhibit transformation; *SOS1* KO significantly enhanced osimertinib-dependent killing in H1975, PC9, and H827 cells after long-term culture (Fig. 3C) consistent with the synergistic effects on survival previously reported for combined EGFR/SOS1 inhibition in short-term cultures^22^. In contrast, the effect of *SOS2* deletion on osimertinib-dependent 3D transformation and survival was dependent on the culture nutrient conditions; in 10% serum *SOS2* KO had a modest effect on osimertinib-dependent inhibition of transformation, but at lower serum levels *SOS2* KO enhanced osimertinib-dependent inhibition of transformation and survival as assessed by both an overall decrease in AUC in all three cell lines and an EC_50_ shift in both H1975 and HCC827 cells (Figs. 3C and S2).

We then assessed the extent to which *SOS2* KO affected the activation of downstream signaling pathways associated with 3D proliferation and survival in whole-cell lysates of 3D cultured spheroids. In H1975 and PC9 cells, *SOS2* KO did not alter ERK phosphorylation as a surrogate of RAF/MEK/ERK signaling in either 10% or 2% serum. In contrast, *SOS2* KO decreased AKT phosphorylation as a surrogate of PI3K/AKT signaling, and this inhibition was more pronounced at low serum conditions (Fig. 3D). These data support previous studies describing the differential preference of SOS2 for promoting EGF-stimulated PI3K/AKT activation in *KRAS*-mutated cells^21, 23^.

Resistance to EGFR-TKIs including osimertinib is most often driven by RTK/RAS/PI3K pathway reactivation^10^ via either tertiary EGFR mutations or enhanced signaling through parallel RTKs including MET, AXL, HER2/3, and FGFR^7–9, 11–16^. Since SOS2 enhanced osimertinib-dependent inhibition of PI3K/AKT signaling, we hypothesized that SOS2 could be an important regulator of RTK/PI3K-dependent osimertinib resistance. MET amplification is one of the most common alternative RTK-dependent EGFR-TKI resistance mechanisms; MET-dependent osimertinib resistance can be modeled by exogenous HGF stimulation^34^. To assess the extent to which SOS2 regulates osimertinib resistance driven by alternate RTKs, we assessed osimertinib dose-dependent inhibition of survival and effector signaling in both 2D (adherent) and 3D spheroid cultured NT and *SOS2* KO H1975 cells (Fig. 4). In H1975 cells cultured in 2D conditions, *SOS2* deletion did not significantly alter the sensitivity of cells to osimertinib (Fig. 4A) or limit the upregulated RAF/MEK/ERK or PI3K/AKT signaling via HGF stimulation compared to NT controls. In contrast, 3D spheroid-cultured *SOS2* KO cells showed enhanced osimertinib-dependent inhibition of survival (Fig. 4A) and PI3K/AKT signaling (Fig. 4B) compared to NT controls. Since *SOS2* KO exclusively to inhibited AKT, but not ERK activation, these data suggest that SOS2 is a critical determinant of RTK/PI3K-dependent osimertinib resistance.

**Fig. 4.**
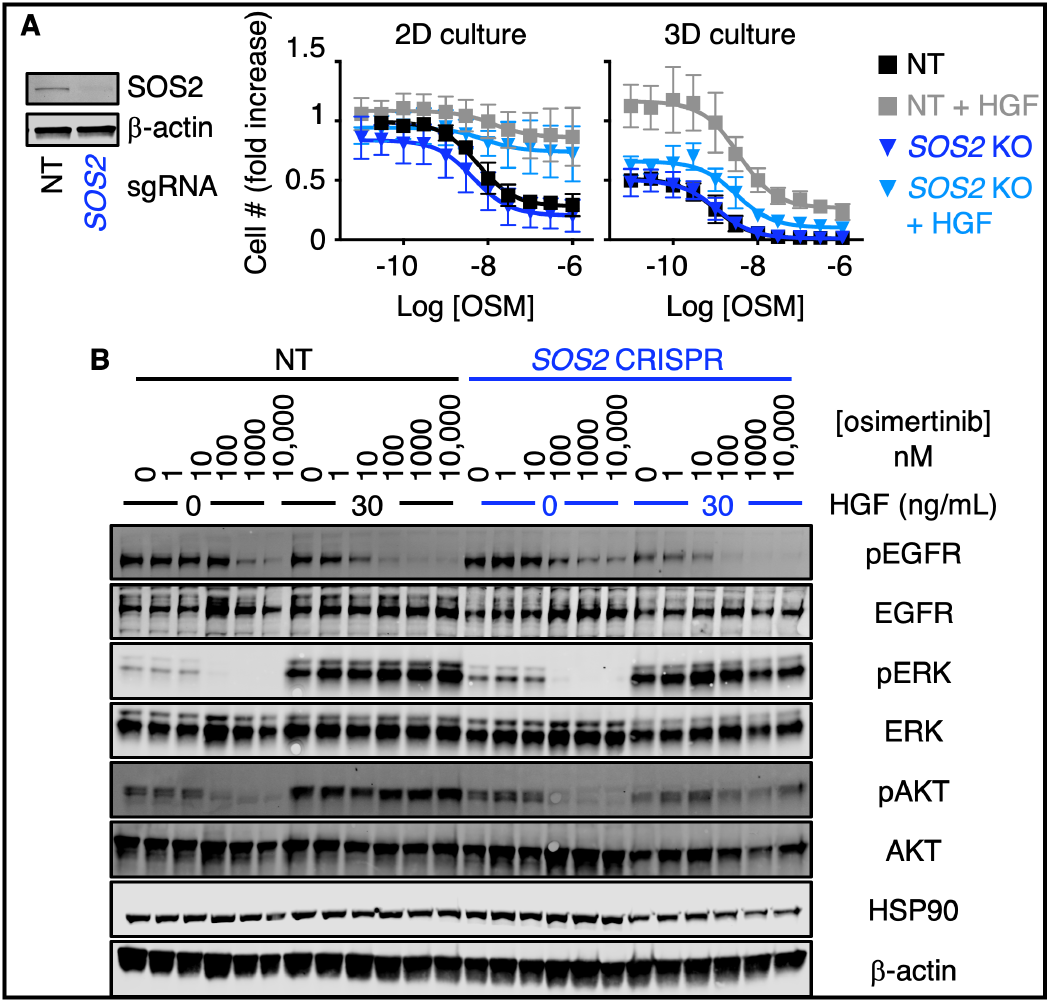
*SOS1* or *SOS2* deletion limits HGF-dependent osimertinib resistance in 3D cultured H1975 cells. (**A**) Western blots of WCLs showing *SOS2* deletion (left) and dose-response curves (right) of *SOS2* KO H1975 cells treated with increasing doses of osimertinib +/-HGF under 2D adherent (top) or 3D spheroid (bottom) culture conditions for 4-days. Dose-response curves are normalized to cell number assessed two hours after plating by CellTitre Glo. (**B**) Western blots of whole cell lysates (WCLs) from NT or *SOS2* KO H1975 cells treated with increasing doses of osimertinib +/-HGF (to bypass EGFR signaling) for 6 hours versus NT controls. Western blots are for pEGFR, EGFR, pERK, ERK, pAKT, AKT, HSP90, and β-actin. Data in A are presented as mean +/-s.d. from *N*=3 independent experiments. Western blots are representative of *N*=3 independent experiments.

To directly assess the extent to which SOS2 regulates the development of acquired resistance to osimertinib, we used an *in-situ* resistance assay (ISRA)^30^ that acts as a cell culture model of a multiple-subject trial to assess resistance to RTK/RAS pathway inhibitors. This hybrid approach combines elements of time-to-progression assays^35, 36^ and cell outgrowth assays^29, 37, 38^ allowing us to monitor the development of de novo osimertinib resistance.

NT and *SOS2* KO H1975, HCC827, PC9, and PC9-TM cells were seeded at low density in the inner 60 wells of multiple 96 well plates and each plate was treated with a single dose (50 – 300 nM) of osimertinib. Wells were fed and scored weekly; wells that reached ≥ 50% confluence were scored as osimertinib resistant and data were plotted as a Kaplan-Meier curve (Fig. 5). In cells treated with lower doses of osimertinib that delayed outgrowth by 1-2 weeks but did not show the prolonged growth arrest necessary to model resistance (50 nM in all cell lines, 150 nM in PC9 cells), *SOS2* KO significantly delayed the outgrowth of drug-treated populations (Fig. 5, dotted lines). In cells treated with doses of osimertinib sufficient to cause prolonged growth arrest and model drug resistance, *SOS2* KO both delayed the outgrowth of osimertinib-resistant cells and reduced the overall frequency of wells able to develop osimertinib resistance (Fig 5, dashed and solid lines). These data suggest that proximal RTK pathway inhibition, achieved here via *SOS2* deletion, may be a strategy to limit osimertinib resistance.

**Fig. 5.**
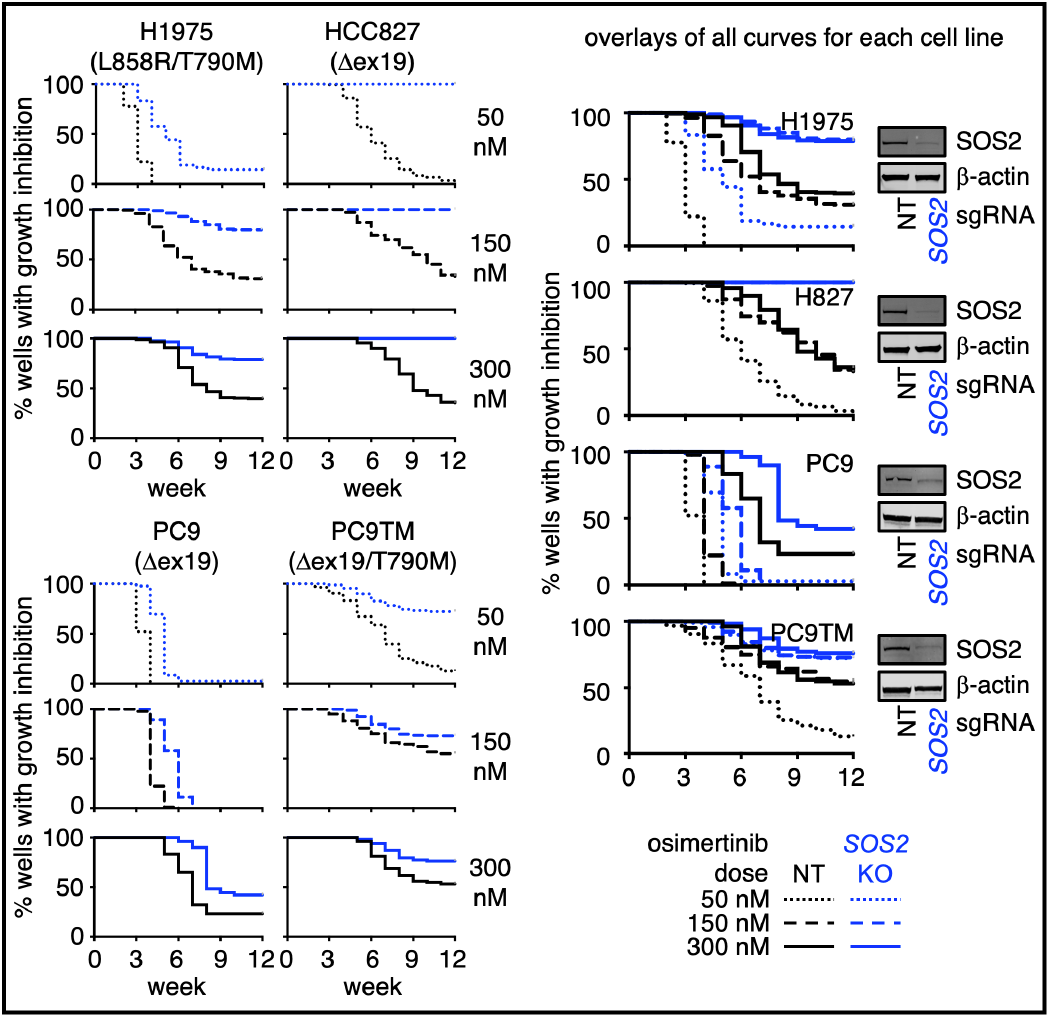
*SOS2 deletion* limits osimertinib resistance in cell culture models. *In situ* resistance assays in NT (black) versus *SOS2* KO (blue) H1975, HCC827, PC9, and PC9-TM cells treated with 50 nM (dotted), 150 nM (dashed), or 300 nM (solid) osimertinib. Separate curves for individual osimertinib doses (left) and overlays of all osimertinib doses (right) are shown for each cell line. Data from *N*=3 independent experiments were combined to generate Kaplan-Meyer curves. Western blots for SOS2 and β-actin (right) are representative of *N*=3 independent experiments.

RTK pathway reactivation^4, 7–16^, often by simultaneous activation of multiple RTKs^30, 39^, represents a common mechanism driving resistance to EGFR-TKIs including osimertinib. RTK-dependent PI3K/AKT activation is a common hallmark of EGFR-TKI resistance^28, 29^, and *SOS2* KO reduced HGF-stimulated PI3K/AKT signaling to inhibit HGF-mediated osimertinib resistance in *EGFR*-mutated cells (Fig. 4). Thus, we hypothesized that the reduced frequency with which *SOS2* KO cultures developed osimertinib resistance in long-term cultures was due to by reduced SOS2-dependent PI3K signaling, and further that *SOS2* KO cultures would become osimertinib resistant via non-RTK dependent mechanisms. To determine whether osimertinib-resistant *SOS2* KO cultures were fundamentally different than NT controls, we expanded 59 NT and 37 *SOS2* KO osimertinib-resistant populations from H1975 cells treated with 150 nM or 300 nM osimertinib for ≥ 6 weeks and assessed whole cell lysates for RTK pathway reactivation (pERK and pAKT) and markers of EMT (E-Cadherin and Vimentin) by Western blotting (Fig. S3). Four isolated *SOS2* KO colonies showed continued SOS2 protein abundance were excluded from our analysis (Fig. S3).

EMT is a dynamic process by which epithelial cells acquire mesenchymal characteristics; the transition from epithelial to mesenchymal phenotypes can be characterized by the loss of E-Cadherin and an increase in Vimentin (Fig. 6A). Epithelial cells are E-Cad^hi^/Vim^lo^ whereas mesenchymal cells are E-Cad^lo^/Vim^hi^. Cells undergoing the epithelial-to-mesenchymal transformation can be either E-Cad^hi^/Vim^hi^ or E-Cad^lo^/Vim^lo^, although E-Cad^hi^/Vim^hi^ is the most well characterized transitional state^17, 40, 41^. This hybrid epithelial/mesenchymal state, also known as partial EMT, is often seen in human cancers^40^ and is associated with resistance to EGFR-TKIs^15, 40–44^. We found that NT osimertinib-resistant H1975 cells predominantly showed a hybrid E/M phenotype where cells were either E-Cad^hi^/Vim^hi^ (42%, black squares) or E-Cad^lo^/Vim^lo^ (20%, open grey circles) (Fig. 6B, black/grey). Further, E-Cad^hi^/Vim^hi^ hybrid E/M populations showed elevated pATK but low pERK (Fig. 6C-D), consistent with previous studies showing that RTK-dependent PI3K/AKT activation is a common hallmark of EGFR-TKI resistance^29^. In contrast, those cells that underwent full EMT (E-Cad^lo^/Vim^hi^, grey circles) showed low pAKT and pERK, suggesting that EMT is an alternative pathway for osimertinib resistance distinct from RTK reactivation.

**Fig. 6.**
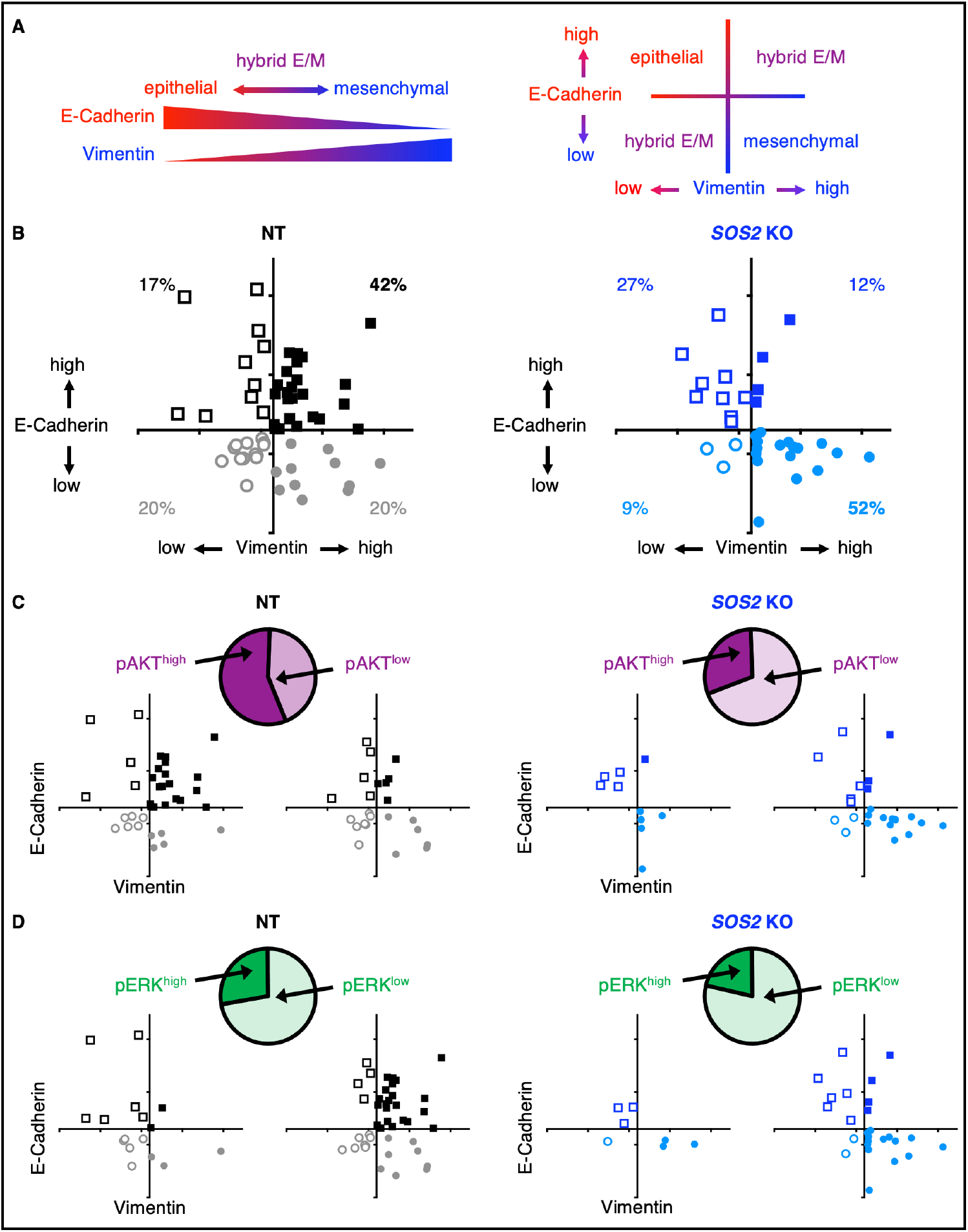
The hybrid E/M phenotype in osimertinib-resistant cells is SOS2-dependent. (**A**) The epithelial-to-mesenchymal transition can be characterized by loss of the epithelial marker E-cadherin and gain of the mesenchymal marker Vimentin; epithelial cells are E-Cad^hi^/Vim^lo^ (open squares) whereas mesenchymal cells are E-Cad^lo^/Vim^hi^ (closed circles). E-Cad^hi^/Vim^hi^ (closed squares) and E-Cad^lo^/Vim^lo^ (open circles) cells are intermediate in this spectrum and constitute a hybrid epithelial/mesenchymal state, also known as ‘partial EMT’. (**B** to **D**) Quantification of E-Cadherin and Vimentin protein abundance from Western blotting experiments in osimertinib-resistant NT and *SOS2* KO H1975 cell populations, either shown as the entire set of cell clones (B) or stratified into pAKT high versus low (C) or pERK high versus low (D) subsets of resistant clones. Each symbol represents an unique osimertinib-resistant colony. NT data are shown in black/grey, SOS2 KO data are shown in light/dark blue. Data are quantified from *N*=3 sets of Western blots.

In contrast to NT controls, the number of osimertinib-resistant populations showing a hybrid E/M phenotype were markedly reduced in *SOS2* KO cells (Fig. 6B). Instead, the few *SOS2* KO populations that were able to become osimertinib-resistant populations primarily underwent full EMT; > 50% of osimertinib-resistant *SOS2* KO populations showed a mesenchymal phenotype (E-Cad^lo^/Vim^hi^) (Fig. 6B, blue) that, similar to NT populations that underwent full EMT, showed low pAKT and pERK indicative of a lack of RTK reactivation (Fig. 6C-D). These data suggest that inhibiting proximal RTK signaling, achieved via *SOS2* deletion, inhibits RTK/AKT-dependent osimertinib resistance but does not inhibit resistance driven by histologic transformation via EMT.

## Discussion

Oncogenic *EGFR* driver mutations occur in 15-30% of lung adenocarcinomas^1–4^. While treatment with the 3^rd^-generation EGFR-TKI osimertinib enhances both progression-free^5^ and overall survival^6^ compared to 1^st^-generation EGFR-TKIs and is the mainstay of therapy for these patients, resistance to osimertinib invariably emerges. Osimertinib resistance is most often driven by reactivation of RAS signaling via activation of multiple parallel RTKs^7–16^ so that single-agent targeting of resistant tumors may be impractical^39^. To prolong the therapeutic window of osimertinib treatment we must identify secondary therapeutic targets whose inhibition either (i) enhances the initial efficacy of osimertinib thereby reducing the overall tumor burden or (ii) inhibits the development of resistant tumor cells by targeting those pathways that drive resistance. Here, we show that the RASGEF SOS2 fulfills each of these criteria: SOS2 modulates the threshold of EGFR signaling to regulate proliferation of *EGFR*-mutated tumors and *SOS2* deletion inhibits RTK/PI3K signaling to block osimertinib resistance driven by oncogenic shift to alternative RTKs.

We previously showed that inhibition of proximal RTK signaling intermediates SOS1 or SHP2 synergistically enhanced the efficacy of osimertinib in short-term (3-4 day) killing assays, but that *SOS2* deletion did not enhance osimertinib efficacy on this timescale^22^. These initial efficacy experiments, similar to most drug-drug synergy studies, were designed to assess secondary targets that would enhance drug-dependent tumor killing but not necessarily inhibition of transforming growth. Further, most *EGFR-*mutated LUAD cell lines grown in 3D require long-term culture (2-3 weeks) to assess for differences in anchorage-independent proliferation^22^. Here, we found that rather than altering transformation under the nutrient-rich conditions used for most experiments, *SOS2* deletion reduced anchorage-independent proliferation when EGFR/RTK stimulation was limiting in both MEFs (Figs. 1-2) and in *EGFR*-mutated LUAD cell lines (Fig. 3). These data extend our original observations that in RTK/RAS mutated cancers, drug-drug synergy should be assessed under 3D culture conditions and suggest that one must also assess the effects of secondary therapeutic targets on multiple timescales to assess both inhibition of 3D spheroid survival (3-4 days) and proliferation (2-3 weeks)^22^.

In addition to enhancing the efficacy of an oncogene-targeted therapy, an ideal co-therapeutic would also delay the development of resistance, thereby enhancing the overall initial window of progression-free survival for the patient receiving treatment. Reactivation of RAS signaling via mutation and/or amplification of multiple parallel RTKs is a common mechanism driving osimertinib resistance^7–16^, and RTK/RAS/PI3K signaling has been hypothesized as a convergent mechanism of EGFR-TKI resistance^29^. SOS2 is critical for RTK-RAS-PI3K signaling in *KRAS*-mutated LUAD cells^21^ and *SOS2* KO reduced PI3K/AKT signaling in osimertinib-treated cells (Fig. 3). Thus, we hypothesized that in addition to enhancing osimertinib efficacy, *SOS2* deletion would delay the onset of osimertinib resistance. To test this hypothesis, we used two distinct models of osimertinib resistance. Using a forced HGF/MET bypass model^34^, *SOS2* deletion re-sensitized HGF-stimulated cells to osimertinib by inhibiting HGF-stimulated PI3K signaling, suggesting that reducing RTK-RAS signaling is sufficient to limit resistance driven by oncogenic shift to an individual RTK. However, this type of ‘forced bypass’ assay does not take into account the evolution cancer cells undergo during long-term selection pressures whereby resistant tumors accrue multiple distinct resistance mechanisms^39^.

To overcome these limitations, we developed an in situ resistance assay that models acquired resistance to RTK/RAS pathway inhibitors in large cohorts of cell populations^30^. Using this assay, we found that *SOS2* deletion reduced the overall frequency with which cultures developed osimertinib resistance (e.g., 68% NT vs. 20% *SOS2* KO in H1975 cells, Fig. 6). Osimertinib resistant populations isolated from ISRAs showed resistance mechanisms similar to patient populations. The majority of resistant populations showed simultaneous hyperactivation of multiple RTKs^30^ and reactivation of PI3K/AKT signaling (Fig. 6), whereas a minority of populations show histologic transformation via EMT (Fig. 6). In contrast, hybrid E/M cells with reactivated RTK/AKT signaling were almost absent from the pool of osimertinib resistant *SOS2* KO cultures. Instead, the few osimertinib-resistant *SOS2* KO cultures that emerged did so primarily by undergoing non-RTK/AKT dependent EMT (Fig. 6). Of note, the overall percentage of osimertinib resistant cultures undergoing full EMT (E-Cad^lo^/Vim^hi^) did not differ between NT [13% of cultures (68% overall resistance x 19% full EMT)] and *SOS2* KO [11% of cultures (20% overall resistance x 51% full EMT)] conditions. These data suggest that targeting proximal RTK signaling has the potential to eliminate the majority of osimertinib resistance, since bypass RTK reactivation and/or tertiary *EGFR* mutations represent the majority of osimertinib-resistant cancers^45^.

In LUAD, RTK/RAS pathway reactivation and ‘oncogene addiction’, or the requirement to maintain elevated RTK/RAS/effector signaling, is not limited to *EGFR*-mutated tumors^4, 28, 46–48^. Indeed, RTK pathway activation is a major resistance mechanism to oncogene-targeted therapies in LUADs with EML-ALK-fusions^48–50^, mutations in other RTKs [*NTRK1*^51^*, ROS1*^52, 53^*, RET*^54^, *MET*^55^, and *HER2* ^56, 57^], or *KRAS* mutations^21, 58–64^. This addiction to RTK/RAS pathway signaling in LUAD suggests that inhibition of proximal RTK signaling is a potential strategy to limit resistance to targeted therapies in a majority of LUADs^4^. The SHP2 phosphatase acts as an adaptor to recruit SOS1 and SOS2 to RTK complexes^65–69^. Thus, in addition to SOS2, SHP2 and SOS1 are RTK signaling intermediates and potential therapeutic targets whose inhibition might limit resistance to RTK/RAS pathway inhibitors in LUAD. In addition to *SOS2* KO, inhibition of proximal RTK signaling via the SHP2 inhibitors RMC-4550 or SHP099 significantly inhibited osimertinib resistance in *EGFR*-mutated LUAD cells^30^. The SOS1 inhibitor BI-3406 significantly inhibited acquired resistance to KRAS^G12C^ inhibitors^70^ or MEK inhibitors^71^ in *KRAS^G^*^12^-mutatated LUAD cells. Based on these data, we propose that inhibition of proximal RTK signaling could be a common mechanism to prevent resistance to targeted therapies in a majority of LUAD.

Our study expands on our previously outlined framework^22^ for preclinical assessment of therapeutic combinations in *EGFR*-mutated cancer cells. Not only do drug-drug synergy experiments need to be performed under 3D culture conditions, but combinations need to be assessed at multiple timeframes to determine the extent to which they enhance drug efficacy (3-4 days), limit oncogenic growth (2-3 weeks), and prevent therapeutic resistance (6-10 weeks, ISRA). Using this framework, our data suggest that SOS2 fulfills the criteria of a secondary therapeutic target that both enhances the efficacy of and reduces resistance to osimertinib in *EGFR*-mutated LUAD.

## Materials and Methods

### Cell culture

Cell lines were cultured at 37°C and 5% CO_2_. HCC827, NCI-H1975, PC9, and PC9-TM cells were maintained in Roswell Park Memorial Institute medium (RPMI), immortalized *Sos1*^f/f^ and *Sos1*^f/f^*Sos2*^-/-^ mouse embryo fibroblasts (MEFs) ^21^ were maintained in Dulbecco’s Modified Eagles Medium (DMEM), each supplemented with 10% fetal bovine serum and 1% penicillin-streptomycin. For 2D signaling experiments, cells were seeded in 10 cm dishes at 1.2 × 10^6^ cells/dish. 24 hours post-plating, cells were treated with inhibitor for 6 hours and then collected for cell lysis and Western blot analysis. For 3D signaling experiments, cells were seeded in 24-well micropatterned AggreWell 400 low-attachment culture plates (Stem Cell # 34415) at 1.2 × 10^6^ cells/well in 2 mL of medium. 24 hours post-plating, half of the media was carefully replaced with fresh media to not disturb the spheroids. At 48 hours, 1 mL media was removed and replaced with 2 x inhibitor. Cells were treated with inhibitor for 6 hours and then collected for cell lysis and Western blot analysis.

### Cell lysis and Western blot analysis

Cells were lysed in RIPA buffer (1% NP-40, 0.1% SDS, 0.1% Na-deoxycholate, 10% glycerol, 0.137 M NaCl, 20 mM Tris pH [8.0], protease (Biotool #B14002) and phosphatase (Biotool #B15002) inhibitor cocktails) for 20 minutes at 4°C and spun at 10,000 RPM for 10 minutes. Clarified lysates were boiled in SDS sample buffer containing 100 mM DTT for 10 minutes prior to Western blotting. Proteins were resolved by sodium dodecyl sulfate-polyacrylamide (Novex precast, ThermoFisher) gel electrophoresis and transferred to nitrocellulose membranes. Western blots were developed by multiplex Western blotting using anti-SOS1 (Santa Cruz sc-256; 1:500), anti-SOS2 (Santa Cruz sc-258; 1:500), anti-β-actin (Sigma AC-15; 1:5,000), anti-pEGFR (Cell Signaling 3777; 1:1000), anti-EGFR (Cell Signaling 4267; 1:1000), anti-pERK1/2 (Cell Signaling 4370; 1:1,000), anti-ERK1/2 (Cell Signaling 4696; 1:1000), anti-pAKT Ser^473^ [Cell Signaling 4060; 1:1000]), anti-AKT (Cell Signaling 2920; 1:1000), anti-HSP90 (Santa Crux sc-7947, 1:1000), anti-α-tubulin (Abcam ab89984; 1:2000), Vimentin (Cell Signaling 5741; 1:1000), and E-Cadherin (Cell Signaling 14472; 1:1000) primary antibodies. Anti-mouse and anti-rabbit secondary antibodies conjugated to IRDye680 or IRDye800 (LI-COR; 1:10,000) were used to probe primary antibodies. Western blot protein bands were detected and quantified using the Odyssey system (LI-COR). For quantification of SOS1 and SOS2 abundance, samples were normalized to either β-actin or HSP90. For quantification of pERK pAKT, E-Cadherin, and Vimentin, samples were normalized to a weighted average of HSP90, β-actin, total ERK1/2, total AKT, and total EGFR ^72^.

### Transformation Studies

MEFs expressing WT or mutant EGFR were seeded in 0.32% Nobel agar at 2 × 10^4^ cells per 35-mm dish to assess anchorage-independent growth. Soft agar colonies were counted 28 days after seeding. For all other cell lines spheroid growth was assessed in ultra-low attachment 96-well round bottomed plates (Corning Costar #7007, S-BIO PrimeSurface #MS-9096UZ, or Nunc Nucleon Sphera microplates ThermoFisher # 174929), cells were seeded at 500 cells per well. Cell number was assessed in parallel plates at 0, 7, 14, and 21 days using CellTiter-Glo® 2.0 reagent.

### sgRNA studies

Cells were infected with lentiviruses (pLentiCRISPRv2 ^73^) expressing Cas9 and either a non-targeting (NT) single guide RNA (sgRNA), a SOS1-targeted sgRNA (SOS1-2), or a SOS2-targeted sgRNA (SOS2-9) as previously described ^21, 22^. Cell lysates were probed for SOS1 or SOS2, and only cell populations (not clones) showing grater that 80% SOS deletion within the overall population were used. Independent infections were used for replicate experiments.

### Production of recombinant lentiviruses

Lentiviruses were produced by co-transfecting MISSION lentiviral packaging mix (Sigma) into 293T cells using Mirus *Trans*IT^®^-Lenti transfection reagent (Mirus Bio # MIR6605) in Opti-MEM (Thermo Scientific #31-985-062). At 48 hours post-transfection, viral supernatants were collected and filtered. Viral supernatants were then either stored at −80°C or used immediately to infect cells in combination with polybrene at 8 μg/mL. 48 hours post-infection, cells were selected in 4 μg/mL Puromycin (Invitrogen). 10-12 days after selection, cells were analyzed for SOS1 and SOS2 protein abundance prior to plating for further studies.

### Inhibitor Studies

For 2D adherent studies cells were seeded at 500-1,000 cells per well in 100 μL in the inner-60 wells of 96-well white-walled culture plates (Perkin Elmer) and allowed to attach for 48 hours prior to drug treatment. Cells were treated with drug for 96 hours (HGF-stimulation studies) or 120 hours (MEFs) prior to assessment of cell viability using CellTiter-Glo® 2.0. For 3D spheroid studies cells were seeded at 500-1,000 cells per well in 100 μL in the inner-60 wells of 96-well ultra-low attachment round bottomed plates (Corning #7007) or Nunc Nucleon Sphera microplates (ThermoFisher # 174929) and allowed to coalesce as spheroids for 48-72 hours prior to drug treatment. For HGF-stimulation studies, cells were treated with osimertinib +/-HGF (30 ng/mL) for 96 hours prior to assessment of cell viability using CellTiter-Glo® 2.0. For transformation studies at different serum concentrations, cells were treated with increasing doses of osimertinib for 7 (PC9) or 21 (H1975, HCC827) days. In all studies parallel plates were assessed for cell viability at the time of drug treatment (day 0) to calculate the fold-change in cell number.

### In situ resistance assays (ISRAs)

ISRAs were performed as previously described ^30^. Briefly, NT and *SOS2* KO cells were seeded at 250 cells/well in the inner 60 wells of replicate 96-well tissue culture plates and allowed to adhere for 24 hours prior to treatment with 50, 150, or 300 nM osimertinib, each plate representing a single drug treatment trial. Plates were fed and wells were scored weekly, with wells reaching >50% confluence scored as resistant. A subset of resistant NT and *SOS2* KO H1975 wells were continuously cultured in osimertinib and expanded prior to whole-cell lysis and assessment by Western blotting.

## Supplementary Materials

**Fig S1** (related to Fig 2). *Sos2* deletion synergizes with EGFR-TKI treatment to inhibit mutant EGFR-driven transformation.

**Fig S2** (related to Fig 3). *SOS2* deletion increases the threshold of EGFR stimulation required for oncogenic transformation.

**Fig S3** (related to Fig 6). The hybrid E/M phenotype in osimertinib-resistant cells is SOS2-dependent.

## Acknowledgments

We thank Udayan Guha for NCI-H1975, HCC827, and PC9 cells and for helpful discussions throughout the project. We thank Julian Downward for PC9-TM cells.

## Author Contributions

PLT and RLK designed the experiments and analyzed the data; PLT and RLK performed most of the experiments; AJL performed Western blots and resistance assays; NES assisted with dose response curves and Wester blots and gave conceptual input throughout the project; BRD assisted with dose response curves and Wester blots; JY assisted with analysis of resistant clones. PLT and RLK wrote the manuscript, NES and BRD edited the manuscript.

## Data and materials availability

All primary data are available on request. All reagents are available from the Kortum laboratory and USUHS via an MTA.

