## Supplemental Figures S12 - S3 for "SOS2 regulates the threshold of mutant *EGFR*-dependent oncogenesis"

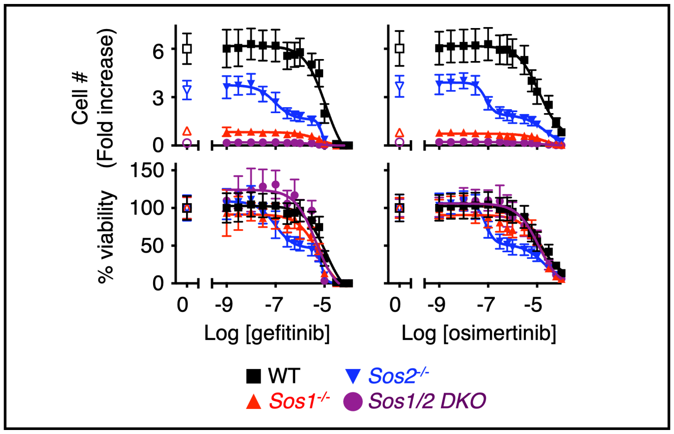


**Fig S1 related to Fig 2**. ***Sos2* deletion synergizes with EGFR-TKI treatment to inhibit mutant EGFR-driven transformation.**

Dose-response curves of WT (black squares), *Sos1*$^{-/-}$ (red triangles), *Sos2*$^{-/-}$ (blue inverted triangles), and *Sos1/2* DKO (purple circles) MEFs expressing EGFR (L858R) from Fig. 1 treated with increasing doses of gefitinib or osimertinib for four days. (Left) Dose-response curves normalized to cell number assessed two hours after plating by CellTitre Glo; (right) dose-response curve where untreated cells for each *Sos1/2* genotype were set to 100%. Data are presented as mean +/- s.d. from three independent experiments.

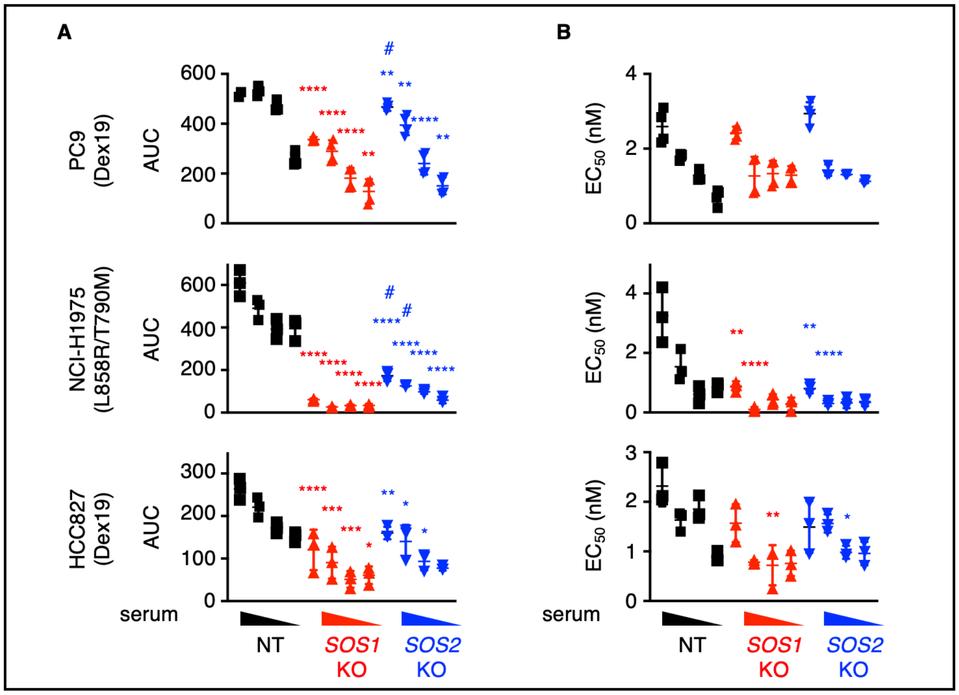


**Fig S2 related to Fig 3. *SOS2* deletion increases the threshold of EGFR stimulation required for oncogenic transformation.**

(**A**) AUC and (**B**) $EC_{50}$ values for osimertinib dose response experiments in 3D spheroid cultured PC9, H1975, or HCC827 cells where *SOS1* or *SOS2* has been deleted using CRISPR/Cas9 versus NT controls shown in Fig. 3C. Cells were treated for 7 (PC9) or 21 (H1975, HCC827) days to allow for transforming growth. Data are presented as mean +/- s.d. from three-four independent experiments.
* $p < 0.05$; ** $p < 0.01$; *** $p < 0.001$; **** $p < 0.0001$ vs. NT controls. # $p < 0.05$ vs. *SOS1* KO.

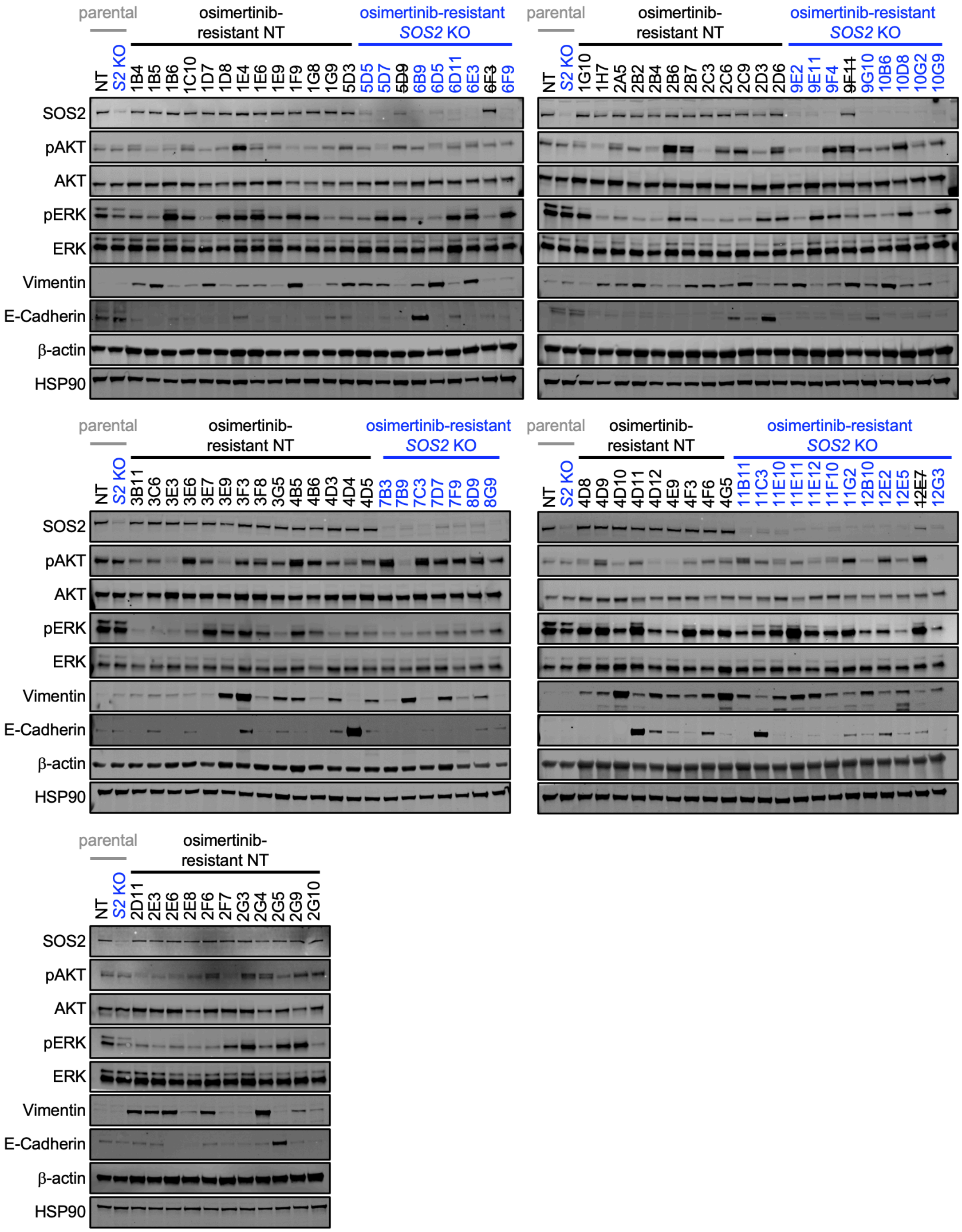


**Fig S3 related to Fig 6. The hybrid E/M phenotype in osimertinib-resistant cells is SOS2-dependent.**

Western blotting for SOS2, pAKT, AKT, pERK, ERK, Vimentin, E-Cadherin, β-actin, and HSP90 in osimertinib-resistant NT and *SOS2* KO H1975 cells.
